# Drug Affinity and Functionalization of Gold Nanoparticles: AFM–SEIRA and SERS Study

**DOI:** 10.64898/2026.05.27.728124

**Authors:** Natalia Piergies, Kamil Raszka, Justyna Wiącek, Magdalena Oćwieja

## Abstract

This study presents the first investigation of the adsorption behaviour of Afatinib on gold nanoparticle (AuNP) monolayers, employing a combination of AFM–SEIRA and SERS techniques. Two types of AuNPs with distinct sizes, synthesized using different reagents, were employed to elucidate the influence of surface type on drug adsorption. The first type of AuNPs was synthesized using sodium borohydride (SB), whereas the second type was obtained using hydroxylamine hydrochloride (HH) as the reducing agent.

AFM–SEIRA revealed that Afatinib interacts to the AuNPs primarily through the quinazoline ring, amide group, and amino moiety, with adsorption geometry strongly dependent on nanoparticle type. Contributions from CH3 and CH2 moieties were also identified, indicating their role in stabilizing the molecule/metal interface. Time–resolved SERS studies demonstrated that the adsorption process is dynamic and involves molecular reorientation, followed by gradual desorption, which is accelerated at physiological temperature (37 °C). Competitive adsorption experiments with phenylboronic acid (PBA) showed that Afatinib exhibits higher affinity toward AuNPs, however, co–adsorption leads to reduced stability of both species on the surface. The results reveal molecular insights into drug/nanoparticle interactions and emphasize the role of surface functionalization in efficient nanocarrier design. This work deepens understanding of adsorption at plasmonic interfaces for biomedical use.

## 1. INTRODUCTION

The development of nanostructured materials for biomedical applications has significantly expanded the possibilities of controlled drug delivery and surface–mediated molecular investigations. In particular, metal nanoparticles (MeNPs) have emerged as promising drug carriers due to their tunable physicochemical properties, high surface–to–volume ratio, and unique plasmonic behaviour. [1,2] At the nanoscale, metals such as silver (Ag), gold (Au) and platinum (Pt) exhibit optical and electromagnetic properties that differ substantially from their bulk counterparts, which makes them attractive platforms for molecular detection, cancer therapy, and nanomedicine–oriented surface engineering.[3–6] Targeted delivery of anticancer agents remains a major challenge in oncology.

The immobilization of drugs onto nanocarrier surfaces may enhance their local concentration at the tumour site and reduce systemic toxicity.[7,8] However, adsorption of bioactive molecules on MeNPs can modify their geometric and electronic structures, potentially affecting therapeutic activity.[9,10] Therefore, detailed characterization of the adsorption geometry, intermolecular interactions, and surface–induced structural rearrangements is essential for the rational design of drug–nanocarrier conjugates. MeNPs support surface plasmon resonance (SPR), a collective oscillation of conduction electrons induced by incident electromagnetic radiation.[11,12] In nanoscale systems, localized surface plasmons (LSPs) generate strongly enhanced and spatially confined electromagnetic fields. These fields constitute the physical basis of surface–enhanced vibrational spectroscopies, namely surface–enhanced Raman spectroscopy (SERS)[13–15] and surface–enhanced infrared absorption (SEIRA).[16,17] The spectral response in surface–enhanced vibrational techniques is strongly dependent on the molecular orientation at the metal interface. According to the surface selection rules,[16,18] only vibrational modes associated with polarizability (in SERS) or dipole moment (in SEIRA) components perpendicular to the metal surface are preferentially enhanced, whereas parallel components are suppressed. These rules provide a powerful framework for deducing adsorption geometry and identifying binding sites of molecules immobilized on metal substrates.[5,6,19–22]

Although SERS and SEIRA provide valuable chemical information, it is limited by diffraction to micrometer–scale spatial resolution.[23] This limitation prevents direct correlation between nanoscale surface morphology and local molecular adsorption behaviour. The combination of atomic force microscopy with infrared spectroscopy (AFM–IR) overcomes this barrier by exploiting the photothermal effect.[24,25] In AFM–IR, pulsed tunable IR radiation induces local thermal expansion proportional to the absorption coefficient of the sample, and this expansion is detected by an AFM probe in contact or tapping mode. As a result, infrared spectra with spatial resolution determined by the AFM tip radius (typically below 20 nm) can be obtained. Importantly, surface enhancement phenomena analogous to SEIRA have been observed in AFM–IR measurements performed on metal nanostructures (AFM–SEIRA effect is visible).[3,5,6,26,27] The local electromagnetic field amplification associated with plasmon excitation may significantly increase the photothermal response, enabling nanoscale mapping of molecular adsorption with enhanced sensitivity. In systems composed of MeNP monolayers, the distribution of AFM–SEIRA signal intensity is expected to correlate with plasmonically active regions, particularly interparticle junctions where electromagnetic coupling is strongest.[3,5] However, the precise contribution of plasmonic effects to AFM–SEIRA signal formation remains an open issue and requires systematic investigations.

In this study, we focus on the adsorption behaviour of Afatinib immobilized on AuNP surfaces. Afatinib is known as an irreversible tyrosine kinase inhibitor (TKI) targeting the ErbB family receptors and is widely used in the treatment of non–small cell lung cancer (NSCLC).[28] Despite its clinical efficacy, Afatinib therapy is associated with dose–dependent adverse effects such as diarrhea, dermatological toxicity, and mucositis.[29] The incorporation of Afatinib into nanostructured delivery systems may improve its bioavailability, reduce required dosage, and minimize systemic toxicity. In this context, understanding how Afatinib interacts with metal nanostructures at the molecular level is crucial for optimizing drug–surface conjugation strategies. Particular attention in this analysis is paid to the influence of the AuNP functionalization strategy with phenylboronic acid (PBA) and Afatinib on the binding affinity of these two molecules and their potential competition for binding sites on the metal surface. Since PBA is capable of specifically binding to sialic acids (SA), which are overexpressed on tumour cell surfaces, including NSCLC,[30,31] such functionalization is expected to enhance the cellular uptake of these nanosystems. By combining AFM–SEIRA with complementary SERS, we aim to elucidate the interaction mechanisms at the Afatinib/PBA/metal interface. Such insight is essential for the rational development of Afatinib–based nanocarrier systems and contributes to a broader understanding of molecule–metal interactions at functionalized nanosurfaces.

## 2. EXPERIMENTAL SECTION

### 2.1. Materials

Hydrogen tetrachloroaurate(III) trihydrate (HAuCl4·3H2O), trisodium citrate dihydrate (Na3C6H5O7·2H2O), hydroxylamine hydrochloride (NH₂OH·HCl), sodium borohydride (NaBH4), sodium chloride (NaCl), poly(allylamine hydrochloride, 56 kDa) (PAH) and sodium hydroxide acid used in the experiments were purchased from Sigma–Aldrich (Merck KGaA, Darmstadt, Germany). All chemicals were used as received, without any additional purification. Ultrapure water (Milli–Q) employed for the preparation of aqueous solutions was produced using a Milli–Q Elix & Simplicity 185 purification system (Millipore SA, Molsheim, France).

### 2.2. Synthesis of AuNPs in the form of hydrosols

Two types of AuNPs were synthesized via chemical reduction using sodium borohydride (SB) and hydroxylamine hydrochloride (HH). For clarity, the nanoparticles are hereafter denoted according to the abbreviations of the respective reducing agents. The preparation procedures for both hydrosols are detailed below.

To obtain SB–AuNPs, 40 mL of a 1.42 mM solution of chloroauric acid was prepared. The solution was then placed under magnetic stirring, and 8 mL of a freshly prepared and cooled 10 mM sodium borohydride solution was added dropwise under continuous stirring. The reaction mixture was further stirred at room temperature for 1 h.

HH–AuNPs were synthesized via chemical reduction carried out at room temperature. Briefly, 0.33 mL of a 1 M sodium hydroxide solution was added dropwise to 90 mL of a 20.3 mM hydroxylamine hydrochloride solution. After 1 min of stirring, 5 mL of a 10 mM chloroauric acid solution was introduced dropwise into the resulting reducing mixture under continuous stirring. Following the completion of the addition, the obtained hydrosol was further stirred for 1 h.

The resulting hydrosols were purified by ultrafiltration using an Amicon 840 stirred cell equipped with a regenerated cellulose membrane (100 kDa molecular weight cutoff) to remove unreacted species and residual impurities. The purification process was continued until the conductivity of the filtrate reached 30 μS·cm⁻¹ and pH 5.8.

The mass concentration of AuNPs in the purified hydrosols was determined based on density measurements of the stock hydrosols and the effluents collected during the washing procedure. Using these values, along with the density of gold (19.3 g cm–3) and the previously described protocol,[32] the concentration of Au in the SB–AuNP and HH–AuNP hydrosols was determined to be 146 mg/L and 173 mg/L, respectively.

### 2.3. Preparation of AuNP monolayers

Well–defined, homogeneous AuNP monolayers with controlled surface coverage were prepared via electrostatically driven deposition, following a previously reported procedure.[3] In brief, thoroughly cleaned calcium fluoride (CaF₂) substrates were immersed in an aqueous PAH hydrosol of concentration equal to 20 mg/L and with an ionic strength of 0.01 M, adjusted by NaCl addition, and a pH of 5.6. The adsorption of the cationic polyelectrolyte onto the CaF₂ surface, driven by electrostatic interactions, was carried out for 20 min, resulting in the formation of a saturated, positively charged PAH layer.[33] Subsequently, the PAH–functionalized CaF₂ substrates were rinsed with Milli–Q water to remove loosely bound polymer chains. The purified, positively charged substrates were then immersed in stock suspensions of SB–AuNP and HH–AuNP (ionic strength 0.01 M, adjusted with sodium chloride). The diffusion–controlled deposition of AuNPs onto the PAH–coated CaF₂ surfaces was allowed to proceed for 5 h. Finally, the samples were gently rinsed with Milli–Q water to to remove excess electrolyte and to prevent sodium chloride crystallization on the surface of the layers, after which the resulting SB–AuNP and HH–AuNP monolayers were dried under ambient conditions.

### 2.4. Samples preparation

Afatinib (purity 99.75%) was obtained from Selleckchem and used as received without further purification. The powdered compound was initially dissolved in Dimethyl sulfoxide (DMSO) to obtain a stock solution with a concentration of 10⁻² M. For AFM–IR and SERS measurements, this stock solution was further diluted with deionized water to 10⁻⁵ M in order to eliminate the influence of DMSO on the recorded spectra. For the SERS measurements, 33 μL of the prepared solution was combined with 33 μL of the stock suspensions of HH–AuNPs (173 mg/L) and SB–AuNPs (146 mg/L), respectively. No additional reagents were introduced. AFM–IR measurements were carried out following the procedure reported previously.[3–5] Briefly, a small droplet of the solution was deposited onto a CaF₂ window and onto the corresponding metal monolayers, after which the samples were allowed to dry under ambient conditions. This preparation strategy enabled direct comparison between AFM–IR spectra of non–adsorbed and surface–bound Afatinib, thereby minimizing the influence of sample preparation on the obtained results.

For the PBA experiments, the compound (purity 99.75%) was purchased from Merck and used without additional purification. The powder was dissolved in deionized water to yield a solution with a concentration of 10⁻³ M. For the SERS measurements, 33 μL of this solution was mixed with 33 μL of the SB–AuNPs stock suspension (146 mg/L), following the same procedure applied for Afatinib. No other reagents were added to the system.

### 2.5. AFM–IR measurements

AFM–IR spectrum of the Afatinib together with the AFM–SEIRA spectra and maps for this drug immobilized on the AuNP monolayers, were acquired using a NanoIR2 system (Anasys Instruments) coupled with a multichip tunable quantum cascade laser (MIRcat–QT, Daylight Solutions) serving as the infrared source. Measurements were performed in tapping mode with commercially available silicon probes coated with a 70 nm gold layer (nominal length 225 μm, nominal width 28 μm), characterized by a resonance frequency of 75 kHz and a spring constant of 3 N/m. For AFM–IR and AFM–SEIRA measurements, 69% and 33%, respectively, of the average QCL output power (0.5 W) were applied, with a pulse duration of 260 ns. Spectra were collected in the range of 1650–1160 cm⁻¹ at a spectral resolution of 2 cm⁻¹. To improve the signal–to–noise ratio (SNR), four spectra were averaged at each measurement point. AFM–SEIRA maps were recorded at a scan rate of 0.5 Hz. The tapping mode setpoint and drive amplitude were set to 6.0 V and 14%, respectively, with 500 points per line. To monitor the sample condition throughout the experiment, AFM topography images were acquired after each series of spectral measurements. Additionally, AFM–SEIRA maps were collected simultaneously with the corresponding topography. This approach confirmed that no thermal damage to the samples occurred under the applied experimental conditions.

### 2.6. RS measurements

The RS spectra of Afatinib and PBA, together with the SERS spectra of NP systems functionalized with Afatinib and PBA using different modification strategies, were recorded using an inVia Renishaw Raman spectrometer. The instrument was equipped with a thermoelectrically cooled CCD detector and a Leica confocal microscope with a 20× objective. Excitation was provided by a 632.8 nm laser line. For each sample, three spectra were collected (one scan per spectrum, 30 s integration time) over the 1800–400 cm⁻¹ spectral range. Reproducibility was assessed by acquiring spectra from three distinct regions of each investigated substrate. No changes in the relative intensities of the characteristic RS and SERS bands were observed, and no signs of sample degradation were detected under the applied experimental conditions.

### 2.7. Data processing

The AFM–IR and AFM–SEIRA data, along with the corresponding 2D intensity maps, were analyzed using Analysis Studio (version 3.14). Three–dimensional intensity maps were constructed with MountainsMap 7.3 software (Digital Surf, France). Prior to analysis, the spectra were smoothed using a Savitzky–Golay filter (third–order polynomial, five data points). Vibrational spectra were processed and visualized using OPUS 7.2 and Origin 22b software. A multipoint baseline correction (five points) was subsequently applied, followed by min–max normalization to prepare the data for interpretation.

## 3. RESULTS AND DISCUSSION

### 3.1. AFM–SEIRA studies

Initially, the AFM–SEIRA approach was used to determine the adsorption geometry of Afatinib on two types of AuNP monolayers with ultra–high spatial resolution. The HH–AuNP monolayer consisted of spherical nanoparticles with an average diameter of 50 nm, whereas the SB–AuNP monolayer was composed of spherical nanoparticles with an average diameter of 20 nm. As shown in Figure 1, which presents the AFM–IR spectrum of non–adsorbed Afatinib together with the AFM–SEIRA spectra recorded after its immobilization on the studied monolayers, the type of AuNP substrate influenced the adsorption mode of the drug on the metal surface. The assignments of the characteristic AFM–IR and AFM–SEIRA bands of Afatinib are summarized in Table 1. The vibrational band assignments were based on conventional and surface–enhanced spectra of quinazoline derivatives,[3,5,6,34] 3–chloro–4–fluoroaniline,[35] tetrahydrofuran and its derivatives.[36]

**Fig. 1.**
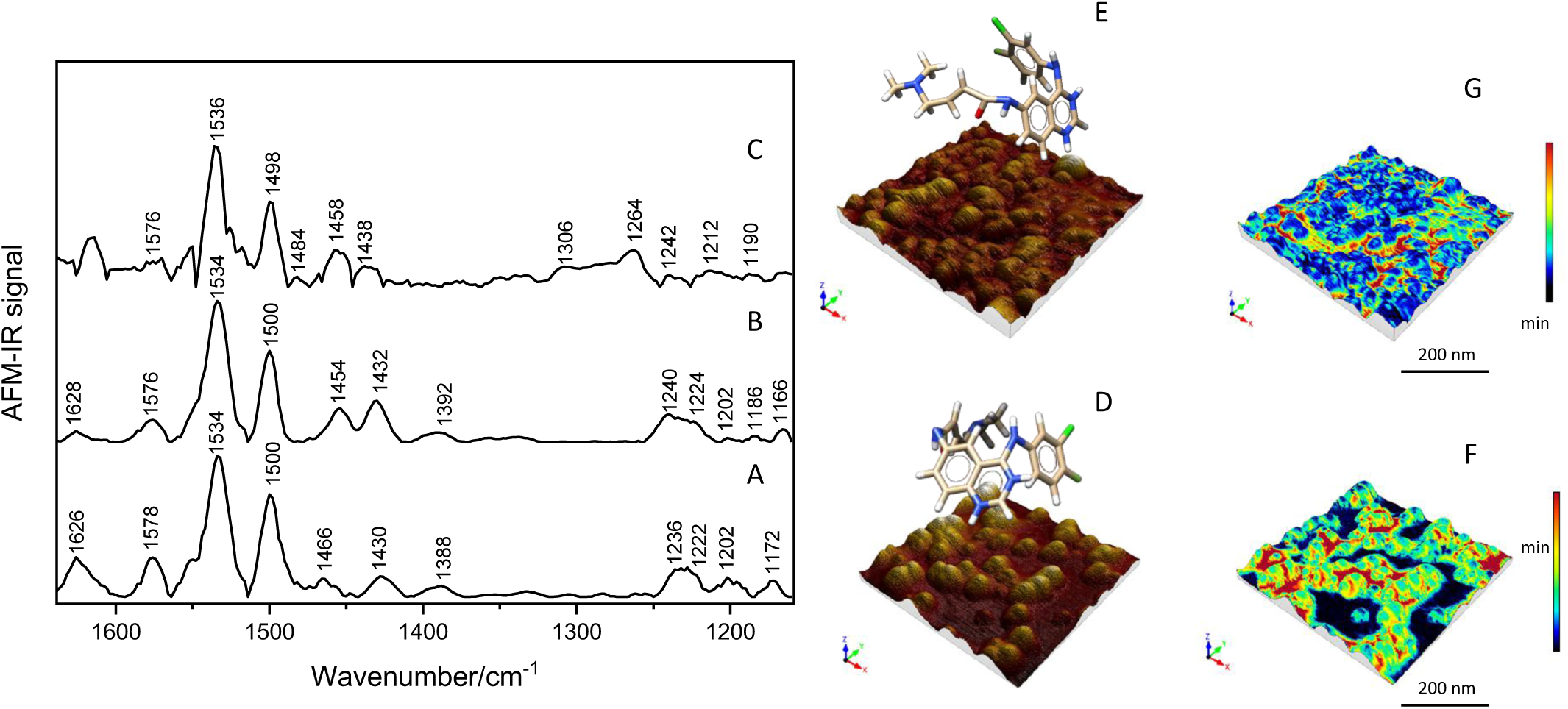
AFM–IR spectrum of Afatinib (A) together with the AFM–SEIRA spectra for this drug immobilized on the HH–AuNPs (B) and SB–AuNPs (C) monolayer, respectively, in the spectral range of 1650 – 1160 cm-1. AFM topographies of Afatinib immobilized on HH–AuNPs (D) and SB–AuNPs (E) monolayers together with the adsorption geometries and corresponding AFM–SEIRA maps showing the spatial distribution of the drug’s spectral signal (F and G, respectively).

**Table 1.**
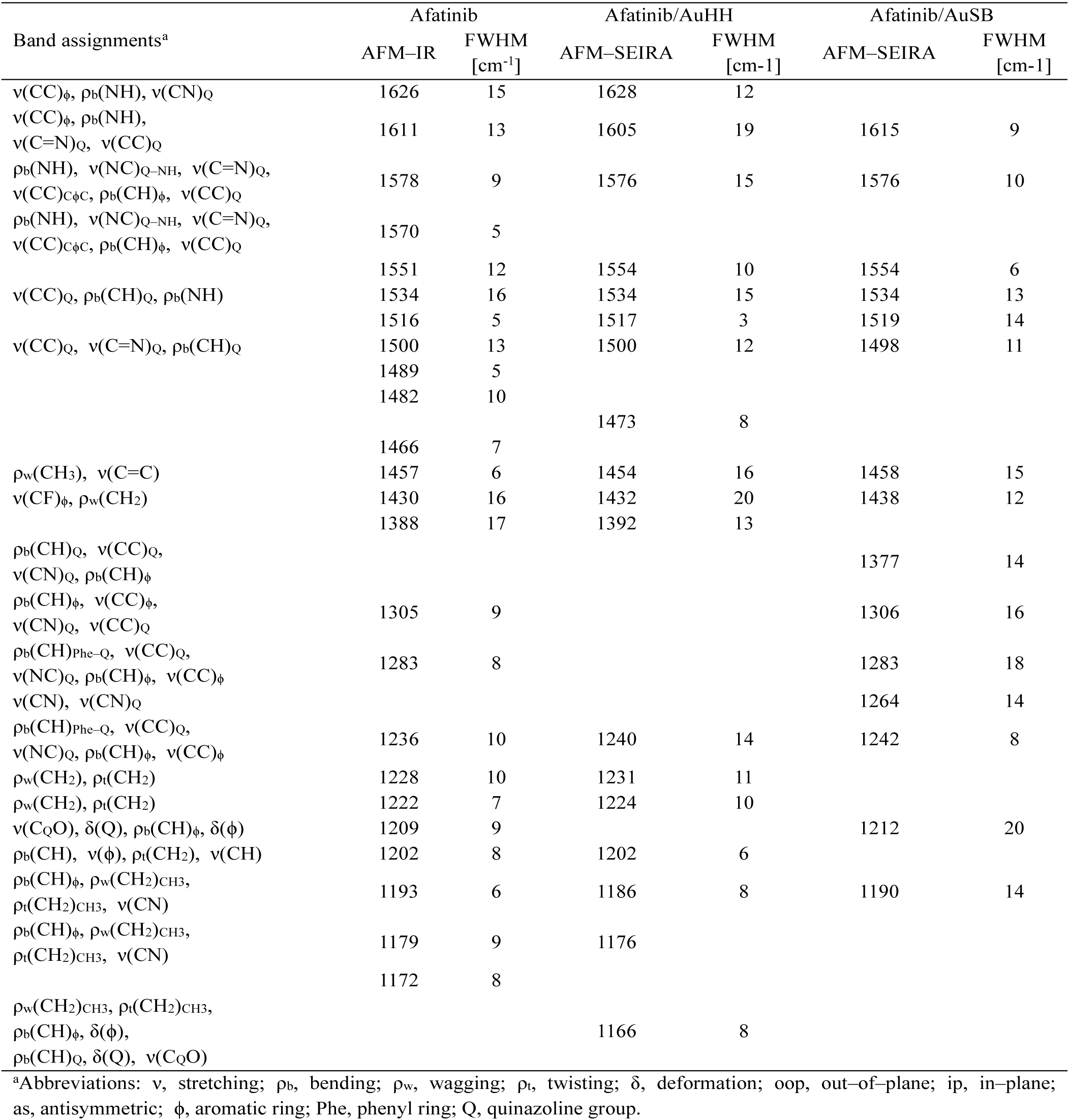
The band assignments together with the wavenumbers (ν) and full width at half maximum (FWHM) for the significant AFM–IR bands of Afatinib and for the AFM–SEIRA bands of this drug after its adsorption on the AgHHNPs and AuSBNPs.[3,5,6,33–35].

| Band assignments<sup>a</sup> | Afatinib |  | Afatinib/AuHH |  | Afatinib/AuSB |  |
| --- | --- | --- | --- | --- | --- | --- |
|  | AFM–IR | FWHM<br>[cm<sup>-1</sup>] | AFM–SEIRA | FWHM<br>[cm<sup>-1</sup>] | AFM–SEIRA | FWHM<br>[cm<sup>-1</sup>] |
| $\nu(\text{CC})_{\phi}$, $\rho_{\text{b}}(\text{NH})$, $\nu(\text{CN})_{\text{Q}}$ | 1626 | 15 | 1628 | 12 | | |
| $\nu(\text{CC})_{\phi}$, $\rho_{\text{b}}(\text{NH})$,<br>$\nu(\text{C}=\text{N})_{\text{Q}}$, $\nu(\text{CC})_{\text{Q}}$ | 1611 | 13 | 1605 | 19 | 1615 | 9 |
| $\rho_{\text{b}}(\text{NH})$, $\nu(\text{NC})_{\text{Q-NH}}$, $\nu(\text{C}=\text{N})_{\text{Q}}$,<br>$\nu(\text{CC})_{\text{C}\phi\text{C}}$, $\rho_{\text{b}}(\text{CH})_{\phi}$, $\nu(\text{CC})_{\text{Q}}$ | 1578 | 9 | 1576 | 15 | 1576 | 10 |
| $\rho_{\text{b}}(\text{NH})$, $\nu(\text{NC})_{\text{Q-NH}}$, $\nu(\text{C}=\text{N})_{\text{Q}}$,<br>$\nu(\text{CC})_{\text{C}\phi\text{C}}$, $\rho_{\text{b}}(\text{CH})_{\phi}$, $\nu(\text{CC})_{\text{Q}}$ | 1570 | 5 | | | | |
|  | 1551 | 12 | 1554 | 10 | 1554 | 6 |
| $\nu(\text{CC})_{\text{Q}}$, $\rho_{\text{b}}(\text{CH})_{\text{Q}}$, $\rho_{\text{b}}(\text{NH})$ | 1534 | 16 | 1534 | 15 | 1534 | 13 |
|  | 1516 | 5 | 1517 | 3 | 1519 | 14 |
| $\nu(\text{CC})_{\text{Q}}$, $\nu(\text{C}=\text{N})_{\text{Q}}$, $\rho_{\text{b}}(\text{CH})_{\text{Q}}$ | 1500 | 13 | 1500 | 12 | 1498 | 11 |
|  | 1489 | 5 |  |  |  |  |
|  | 1482 | 10 |  |  |  |  |
|  |  |  | 1473 | 8 |  |  |
|  | 1466 | 7 |  |  |  |  |
| $\rho_{\text{w}}(\text{CH}_3)$, $\nu(\text{C}=\text{C})$ | 1457 | 6 | 1454 | 16 | 1458 | 15 |
| $\nu(\text{CF})_{\phi}$, $\rho_{\text{w}}(\text{CH}_2)$ | 1430 | 16 | 1432 | 20 | 1438 | 12 |
|  | 1388 | 17 | 1392 | 13 |  |  |
| $\rho_{\text{b}}(\text{CH})_{\text{Q}}$, $\nu(\text{CC})_{\text{Q}}$,<br>$\nu(\text{CN})_{\text{Q}}$, $\rho_{\text{b}}(\text{CH})_{\phi}$ | | | | | 1377 | 14 |
| $\rho_{\text{b}}(\text{CH})_{\phi}$, $\nu(\text{CC})_{\phi}$,<br>$\nu(\text{CN})_{\text{Q}}$, $\nu(\text{CC})_{\text{Q}}$ | 1305 | 9 | | | 1306 | 16 |
| $\rho_{\text{b}}(\text{CH})_{\text{Phe-Q}}$, $\nu(\text{CC})_{\text{Q}}$,<br>$\nu(\text{NC})_{\text{Q}}$, $\rho_{\text{b}}(\text{CH})_{\phi}$, $\nu(\text{CC})_{\phi}$ | 1283 | 8 | | | 1283 | 18 |
| $\nu(\text{CN})$, $\nu(\text{CN})_{\text{Q}}$ | | | | | 1264 | 14 |
| $\rho_{\text{b}}(\text{CH})_{\text{Phe-Q}}$, $\nu(\text{CC})_{\text{Q}}$,<br>$\nu(\text{NC})_{\text{Q}}$, $\rho_{\text{b}}(\text{CH})_{\phi}$, $\nu(\text{CC})_{\phi}$ | 1236 | 10 | 1240 | 14 | 1242 | 8 |
| $\rho_{\text{w}}(\text{CH}_2)$, $\rho_{\text{t}}(\text{CH}_2)$ | 1228 | 10 | 1231 | 11 | | |
| $\rho_{\text{w}}(\text{CH}_2)$, $\rho_{\text{t}}(\text{CH}_2)$ | 1222 | 7 | 1224 | 10 | | |
| $\nu(\text{C}_\text{QO})$, $\delta(\text{Q})$, $\rho_{\text{b}}(\text{CH})_{\phi}$, $\delta(\phi)$ | 1209 | 9 | | | 1212 | 20 |
| $\rho_{\text{b}}(\text{CH})$, $\nu(\phi)$, $\rho_{\text{t}}(\text{CH}_2)$, $\nu(\text{CH})$ | 1202 | 8 | 1202 | 6 | | |
| $\rho_{\text{b}}(\text{CH})_{\phi}$, $\rho_{\text{w}}(\text{CH}_2)_{\text{CH}_3}$,<br>$\rho_{\text{t}}(\text{CH}_2)_{\text{CH}_3}$, $\nu(\text{CN})$ | 1193 | 6 | 1186 | 8 | 1190 | 14 |
| $\rho_{\text{b}}(\text{CH})_{\phi}$, $\rho_{\text{w}}(\text{CH}_2)_{\text{CH}_3}$,<br>$\rho_{\text{t}}(\text{CH}_2)_{\text{CH}_3}$, $\nu(\text{CN})$ | 1179 | 9 | 1176 | | | |
|  | 1172 | 8 |  |  |  |  |
| $\rho_{\text{w}}(\text{CH}_2)_{\text{CH}_3}$, $\rho_{\text{t}}(\text{CH}_2)_{\text{CH}_3}$,<br>$\rho_{\text{b}}(\text{CH})_{\phi}$, $\delta(\phi)$,<br>$\rho_{\text{b}}(\text{CH})_{\text{Q}}$, $\delta(\text{Q})$, $\nu(\text{C}_\text{QO})$ | | | 1166 | 8 | | |

<sup>a</sup>Abbreviations: $\nu$ , stretching; $\rho_{\text{b}}$ , bending; $\rho_{\text{w}}$ , wagging; $\rho_{\text{t}}$ , twisting; $\delta$ , deformation; oop, out-of-plane; ip, in-plane; as, antisymmetric; $\phi$ , aromatic ring; Phe, phenyl ring; Q, quinazoline group.

As the applicability of SEIRA surface selection rules to AFM–SEIRA data has been previously demonstrated,[5,6] these rules were used to interpret the recorded spectra. For both monolayers, the surface–enhanced spectra are dominated by bands associated with vibrations of the quinazoline moiety. These appear at ∼1576 cm⁻¹, 1498 cm⁻¹, and 1236 cm^-1^ corresponding to the ν(NC)_Q–NH_/ν(C=N)_Q_/ν(CC)_CϕC_/ρ_b_(CH)_ϕ_, ν(CC)_Q_, ν(CC)_Q_/ν(C=N)_Q_/ρ_b_(CH)_Q_, and ρ_b_(CH)_Phe–Q_/ν(CC)_Q_/ν(NC)_Q_ modes, respectively. A particularly strong interaction involving the C–N bond of the quinazoline ring is observed for the SB–AuNP monolayer, as indicated by the intense band at 1264 cm⁻¹, which is present only in the AFM–SEIRA spectrum of this surface.

The participation of the 3–chloro–4–fluoroaniline moiety in the Afatinib/AuNPs interaction is evidenced by the bands observed at ∼1628 cm⁻¹, ∼1432 cm⁻¹, and ∼1186 cm⁻¹, assigned to the ν(CC)_ϕ_, ν(CF)_ϕ_, and ρ_b_(CH)_ϕ_ vibrations, respectively. However, it should be noted that the bands associated with the C–F bond modes exhibit stronger enhancement for the HH–AuNP monolayers than for the SB–AuNP monolayers, indicating stronger interaction with the metal surface in the former case. This conclusion is corroborated by the band shift (Δν) and broadening (ΔFWHM) of this spectral feature relative to the corresponding band in the AFM–IR spectrum of the drug prior to adsorption (Δν = 2 cm⁻¹ and ΔFWHM = 4 cm⁻¹). Also, the aliphatic moieties of Afatinib take part in the studied molecular/metal interaction. The NH group interaction with both monolayers is confirmed by the high–intensity band at ∼1536 cm⁻¹ attributed to the ρ_b_(NH) modes. The CH_3_ from dimethylamine and CH_2_ from but–2–enamide moieties of Afatinib indicate interaction with the NPs. Namely, band at ∼1454 cm^-1^, ∼1228 cm^-1^, ∼1224 cm^-1^, and ∼1166 cm^-1^ responsible for the ρ_w_(CH_3_), ρ_w_(CH_2_)/ρ_t_(CH_2_), ρ_w_(CH_2_)/ρ_t_(CH_2_), and ρ_w_(CH_2_)_CH3_/ρ_t_(CH_2_)_CH3_. These low–and medium–intensity bands indicate that the CH₂ and CH₃ groups are located in close proximity to the metal monolayers. The suggested adsorption geometry of Afatinib on the HH–AuNP and SB–AuNP monolayers is illustrated in Figures 1D and 1E, respectively.

The AFM–SEIRA technique, in addition to providing information on the nanoscale spatial orientation of molecules on MeNP monolayers, also reveals a strongly enhanced spectral signal distribution originating from internanoparticle regions, as well as from the gap between the gold–coated tip and the nanoparticles. Figures 1F and 1G present AFM–SEIRA maps illustrating the spatial distribution of the spectral signal across the HH–AuNP and SB–AuNP monolayers, respectively. The visualized enhancement pattern clearly indicates that the most intense signals are localized in the internanoparticle gaps, confirming that the strongest plasmonic coupling occurs between neighboring nanoparticles rather than between the tip and the nanoparticles.

### 3.2. SERS studies

SERS technique was employed to characterize the temporal changes of the Afatinib/AuNP interaction during adsorption, as well as its dependence on temperature, specifically at room temperature (RT) and 37 °C. Moreover, competitive adsorption between two components, namely: Afatinib and PBA, on the AuNP surface was studied.

Figure 2 shows the RS spectrum of non–adsorbed Afatinib together with the SERS spectra of the drug after adsorption on SB–AuNPs, recorded within 30 min, 24 h, and 48 h of incubation at RT. Table 2 presents the band assignments based on the conventional RS and SERS spectra for quinazoline derivatives,[4,5,34,37] amide bonds,[38,39] and dimethylamine derivative.[40] The SERS spectrum collected 30 min after adsorption indicates that the interaction between Afatinib and the AuNP surface is predominantly mediated by the quinazoline moiety. This conclusion is supported by the pronounced enhancement of the bands located at 1623 cm^-1^, 1397 cm^-1^, 1382 cm^-1^, 1295 cm^-1^, 690 cm^-1^ which are attributed to the ν(CC)_Q_/ν(CN)_Q_, ν(C=N)_Q_/ν(CC)_Q_/ρ_b_(CH)_Q_, ν(CC)_Q_/ρ_b_(CH)_Q_, ν(CC)_Q_/ρ_b_(CH)_Q_, ν_i.p_.(Q) modes, accordingly. In addition, vibrational modes of the quinazoline ring contribute to the band observed at 778 cm⁻¹, assigned to the ν(Q) stretching vibration. Comparison of these SERS bands with the corresponding features in the RS spectrum reveals noticeable band broadening (∼4 cm⁻¹) and position shifts (∼5 cm⁻¹; see Table 2). These spectral changes indicate a relatively strong interaction between the quinazoline fragment and the SB–AuNPs. Furthermore, the dominant enhancement of in–plane vibrational modes suggests that the quinazoline ring adopts an orientation close to perpendicular with respect to the metal surface. This interpretation is in agreement with the SERS surface selection rules[16,18] discussed in the Introduction. Such an adsorption geometry implies that the N=C bond of the quinazoline unit is directed toward the SB–AuNPs, facilitating the interaction with the studied metal substrate.

**Table 2.**
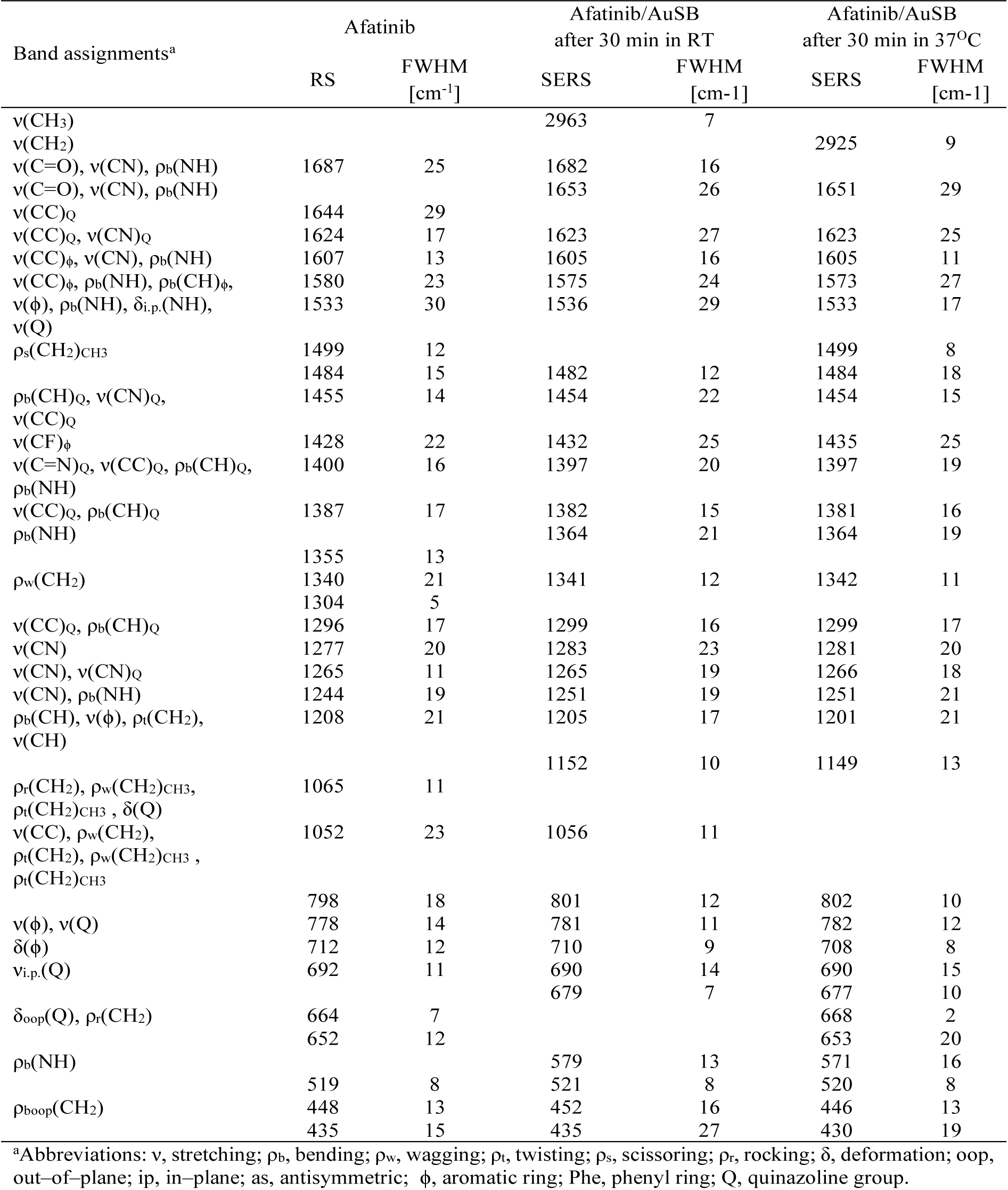
The band assignments together with the wavenumbers (ν) and full width at half maximum (FWHM) for the significant RS bands of Afatinib and for the SERS bands of this drug after its adsorption on the AuSBNPs.[4,5,33,36–39].

Additionally, the involvement of the 3–chloro–4–fluoroaniline moiety in the Afatinib/SB–AuNPs interaction can be observed. The bands at 1605 cm^-1^ [ν(CC)_ϕ_], 1575 cm^-1^ [ν(CC)_ϕ_], 1205 cm^-1^ [ν(ϕ)], 781 cm^-1^ [ν(ϕ)], and 710 [δ(ϕ)] display weaker or nearly unchanged intensities in the SERS spectrum compared with the corresponding features in the RS spectrum. Such a spectral pattern suggests that this aromatic fragment is located in the vicinity of the AuSBNP surface, although it is not directly involved in the adsorption. According to the SERS surface selection rules, the relatively low enhancement of these vibrational modes, together with the limited band broadening, indicates that the phenyl ring of the 3–chloro–4–fluoroaniline unit interacts more weakly with the SB–AuNPs than the quinazoline fragment. Additionally, the low intensity of bands assigned to the C–F and C–Cl stretching vibrations (see Table 2) suggests that these substituents are oriented away from the metal surface and therefore do not significantly contribute to the adsorption process.

Similarly to the observations obtained from AFM–SEIRA measurements, the SERS spectra also provide evidence of the interaction between the but–2–enamide fragment and the SB–AuNPs. The intense band observed at 1653 cm⁻¹ is associated with the ν(C=O)/ν(CN)/ρ_b_(NH) vibrational modes, indicating that the amide bond interacts strongly with the metal surface. The involvement of the amino groups is also clearly reflected in the SERS spectrum. The strongly enhanced bands at 1536 cm^-1^, 1283 cm^-1^, 1265 cm^-1^, and 1251 cm^-1^ are assigned to the ρ_b_(NH)/δ_i.p._(NH), ν(CN), ν(CN), ν(CN), and ν(CN)/ρ_b_(NH) vibrations, respectively. This spectral pattern, together with the observed band shifts and broadening relative to the corresponding RS spectral features (see Table 2), suggests that the amino group originating from the but–2–enamide fragment, as well as the amino group located between the quinazoline and 3–chloro–4–fluoroaniline moieties, interacts directly with the SB–AuNPs surface.

The SERS technique also enables monitoring of the temporal stability of the interaction between the investigated drug and SB–AuNPs. Figure 3C and D illustrate the spectral signals of Afatinib immobilized on the AuNP surface after 24 h and 48 h of adsorption, respectively. As can be observed, after 24 h of adsorption the bands at 1266 cm⁻¹ [ν(CN)] and 1251 cm⁻¹ [ν(CN)/ρ_b_(NH)] exhibit increased intensity. This behaviour suggests a reorientation of the drug molecules on the SB–AuNPs, in which the amino groups begin to interact more strongly with the metallic substrate. Such reorientation likely leads to a gradual weakening of the overall adsorption, resulting in slow drug desorption. This process becomes apparent after 48 h, as indicated by the decreased signal–to–noise ratio observed in the SERS spectrum (Figure 3D). The suggested adsorption behaviour of the drug on the SB–AuNPs is shown in Figure 2E and 2F.

**Fig. 2.**
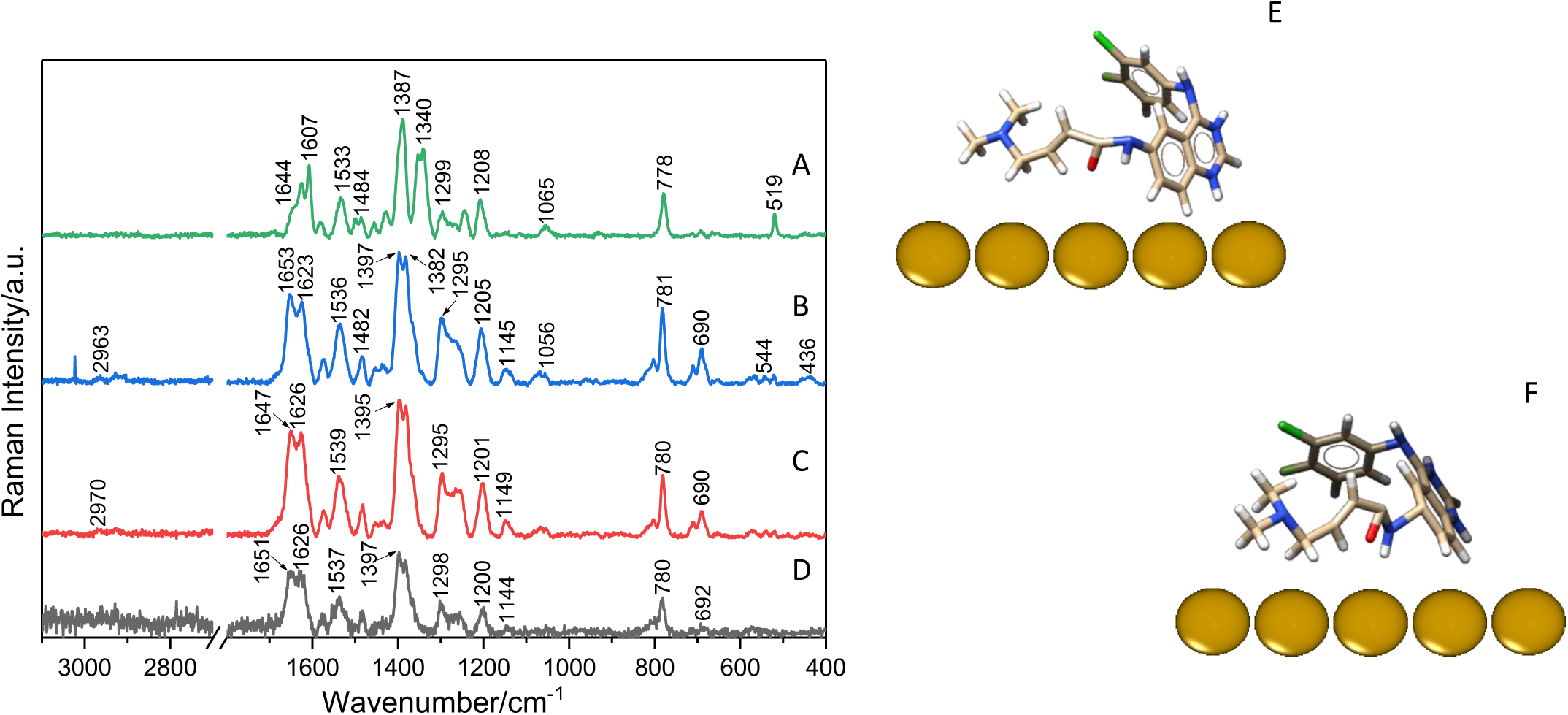
RS spectrum of Afatinib (A) together with the SERS spectra for this drug immobilized on the SB–AuNPs after 30 min (B), 24 h (C), and 48 h (D) adsorption time in the room temperature, respectively, in the spectral range of 3100–400cm-1, along with the evolution of drug reorganization on the NPs after 30 min (E) and 24 h (F).

**Fig. 3.**
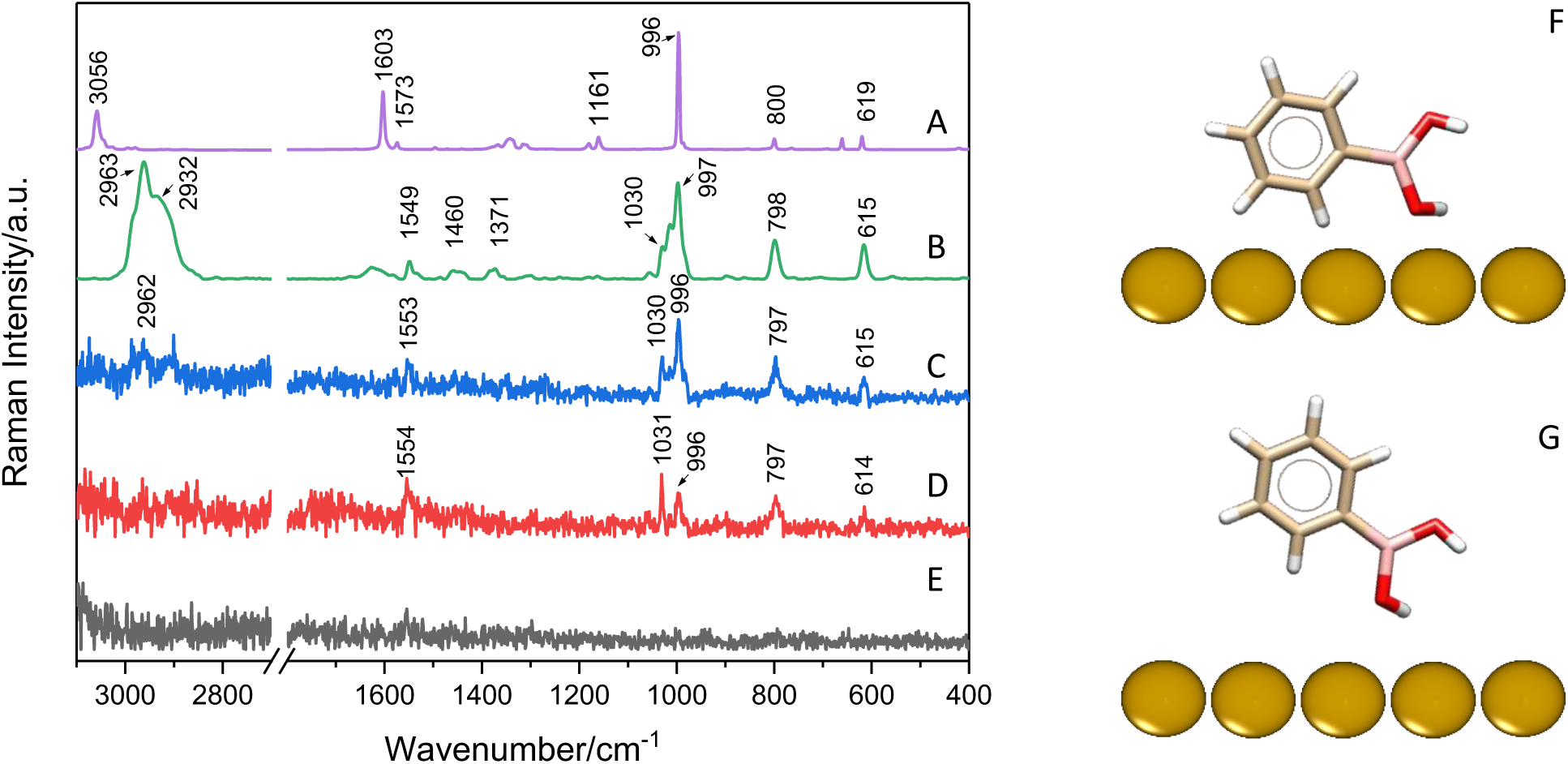
Raman spectrum of PBA (A) and SERS spectra of PBA immobilized on SB–AuNPs after 30 min (B) and 24 h (C) at room temperature, as well as after 30 min (D) and 24 h (E) at 37 °C adsorption time, in the spectral range of 3100–400cm-1, together with the suggested adsorption mode of PBA after 30 min in RT (F) and 37 °C (G), respectively.

When considering the influence of temperature on the Afatinib/SB–AuNPs interaction, it can be observed that after 30 minutes the adsorption, behaviour of the drug at 37 °C is nearly identical to that observed at RT (see Figures S1 and 3B). The most intense spectral bands are associated with vibrations of the quinazoline moiety, the amide bond, and amino groups. In addition, the 3–chloro–4–fluoroaniline fragment also participates in the adsorption process on the NP surface (see Table 2 for assignments). However, a slightly lower intensity of the band at 1650 cm⁻¹, attributed to the ν(C=O)/ν(CN)/ρ_b_(NH) vibrations, suggests a weaker interaction of the amide bond with the SB–AuNPs. At the same time, the increased intensity of the bands at 1266 cm⁻¹ and 1251 cm⁻¹ (see Table 2 for assignments) in the SERS spectrum recorded at 37 °C after 30 min of adsorption, compared with the corresponding spectrum collected at RT, indicates a stronger involvement of the amino group in the adsorption process. This interaction, similarly to that observed at RT, appears to represent the initial stage of drug reorientation on the NP surface, ultimately leading to its desorption. This interpretation is supported by the observed decrease in intensity of bands associated with the amide group (1650 cm⁻¹) and the quinazoline moiety (1397 cm⁻¹, 781 cm⁻¹, 690 cm⁻¹), accompanied by an increase in the intensity of the spectral features at 1266 cm⁻¹ and 1251 cm⁻¹. Finally, after 24 h of adsorption, desorption of Afatinib from the SB–AuNPs is observed (see Figure S1).

The subsequent part of the study focuses on the adsorption mode of PBA molecules. Similarly to the case of Afatinib, the influence of adsorption time and temperature on the adsorption behaviour was also investigated. Figure 4 presents the RS and SERS spectra of non–adsorbed PBA and after its adsorption on SB–AuNPs at RT and 37 °C, respectively. Table 3 present the performed band assignments based on the RS and SERS spectra of PBA derivatives.[41–44] At RT, the most intense bands observed in the SERS spectrum are attributed to in–plane vibrational modes of PBA. In particular, the spectral features at 1576 cm⁻¹, 1442 cm⁻¹, 1031 cm⁻¹, and 997 cm⁻¹, assigned to ν(CC)ϕ, ν(CC)/δ(HCC), ν(CC), and δ_i.p._(ϕ) vibrations, respectively, exhibit strong enhancement compared with the corresponding RS spectrum. Such spectral behaviour indicates a perpendicular orientation of the PBA molecule relative to the SB–AuNPs surface. Furthermore, an enhancement of the bands at 1014 cm⁻¹ and 798 cm⁻¹, attributed to ν(BO) and ν(BC)/ν(OB) vibrations, respectively, is observed together with noticeable band broadening and shifts in their positions compared with the corresponding RS spectrum (see Table 3). This spectral pattern suggests that PBA adsorbs on the SB–AuNPs surface *via* a perpendicular phenyl ring orientation, with the BO bond directed toward the nanoparticle surface. However, this interaction appears to be unstable. After 24 h, the disappearance of the characteristic PBA spectral bands, accompanied by a decreased signal–to–noise ratio, suggests gradual molecular desorption from the surface. The reduced intensity of bands associated with the BO group indicates that molecular reorientation occurs, leading to a weakening of the BO/SB–AuNPs interaction and ultimately resulting in desorption. This desorption process proceeds even more rapidly at 37 °C. After only 30 min, the recorded spectral pattern resembles that observed for the SERS spectrum collected after 24 h of adsorption at RT. After 24 h at 37 °C, no detectable spectral signal of PBA is observed on the investigated metal surface. Figures 4F and 4G illustrate the proposed PBA adsorption pattern on the SB–AuNPs after 30 minutes at RT and 37 °C, respectively.

**Fig. 4.**
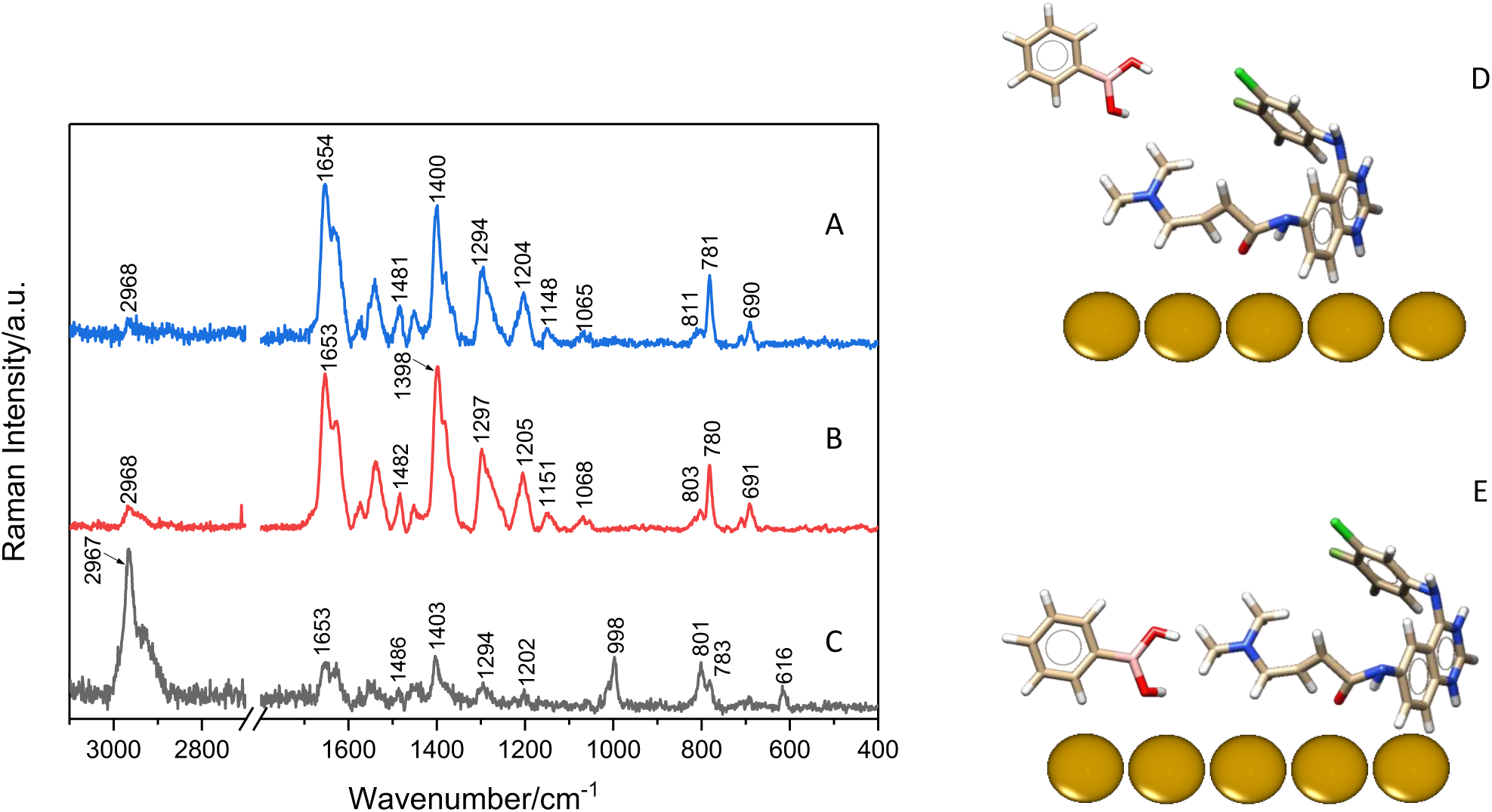
SERS spectra of three systems: AuNPs+(Afatinib+PBA) (A), AuNPs+Afatinib+PBA (B), and AuNPs+PBA+Afatinib (C) recorded after 30 minutes of adsorption at RT, in the spectral range of 3100–400cm-1. Order of the listed components reflects the sequence of molecular attachment. The suggested adsorption behaviour of Afatinib and PBA for the AuNPs+(Afatinib+PBA) and AuNPs+Afatinib+PBA systems (D) and for the AuNPs+PBA+Afatinib system (E).

**Table 3.** The band assignments together with the wavenumbers (ν) and full width at half maximum (FWHM) for the significant RS bands of PBA and for the SERS bands of this drug after its adsorption on the AuSBNPs.[40–43].

| Band assignments<sup>a</sup> | PBA |  | Afatinib/AuSB<br>after 30 min in RT |  |
| --- | --- | --- | --- | --- |
|  | RS | FWHM<br>[cm<sup>-1</sup>] | SERS | FWHM<br>[cm<sup>-1</sup>] |
| $\nu(\text{CC})$, $\delta(\text{HCC})$ | | | 1621 | 65 |
| $\nu(\text{CC})$ | 1603 | 7 | | |
| $\nu(\text{CC})$ | 1574 | 6 | 1576 | 40 |
| $\nu(\text{CC})$ | | | 1549 | 11 |
|  |  |  | 1533 | 19 |
| $\nu(\text{CC})$, $\delta(\text{HCC})$ | 1495 | 6 | | |
|  | 1471 | 3 |  |  |
|  |  |  | 1460 | 14 |
| $\nu(\text{CC})$, $\delta(\text{HCC})$ | | | 1442 | 23 |
|  | 1377 | 16 | 1376 | 26 |
| $\nu(\text{CC})$, $\delta(\text{HOB})$, $\nu(\text{CB})$ | 1367 | 9 | | |
| $\nu(\text{CC})$ | 1181 | 8 | | |
| $\nu(\text{CC})$, (HCC) | 1160 | 7 | | |
|  |  |  | 1054 | 21 |
| $\nu(\text{CC})$ | | | 1031 | 8 |
| $\nu(\text{BO})$ | | | 1014 | 17 |
| $\delta_{\text{i.p.}}(\phi)$ | 996 | 5 | 997 | 13 |
| $\nu(\text{BO})$ | 985 | 4 | 983 | 11 |
|  |  |  | 896 | 14 |
| $\delta(\text{CCC})$, $\nu(\text{BC})$, $\nu(\text{OB})$ | 799 | 5 | 798 | 17 |
| $\delta(\text{CCC})$, $\nu(\text{BC})$, $\nu(\text{CC})$ | 764 | 9 | 770 | 25 |
|  |  |  | 708 | 30 |
| $\rho_{\text{t}}(\text{CCCC})$, $\rho_{\text{t}}(\text{HCCC})$, $\rho_{\text{t}}(\text{OBCC})$ | 693 | 11 | | |
| $\rho_{\text{t}}(\text{CCCH})$, $\rho_{\text{t}}(\text{CCCC})$, $\delta_{\text{oop}}(\text{OH})$ | 660 | 4 | | |
|  | 619 | 5 | 615 | 15 |
| $\rho_{\text{t}}(\text{HOBC})$ | | | 556 | 18 |

<sup>a</sup>Abbreviations: $\nu$ , stretching; $\rho_{\text{t}}$ , twisting; $\delta$ , deformation; oop, out-of-plane; ip, in-plane.

The SERS investigations performed for Afatinib and PBA revealed notable differences in the stability of their interactions with SB–AuNPs. Therefore, the final stage of this study aimed to examine how these two compounds compete for adsorption sites on the metal surface. Figure 4 presents the SERS spectra of three systems: AuNPs+(Afatinib+PBA), AuNPs+Afatinib+PBA, and AuNPs+PBA+Afatinib, representing different sequences of molecular attachment. The spectra shown in Figures 4A and 4B exhibit nearly identical spectral profiles. In both cases, only bands characteristic of Afatinib are observed. The most prominent features appear at ∼1654 cm⁻¹ [ν(C=O)/ν(CN)/ρ_b_(NH)], ∼1400 cm⁻¹ [ν(C=N)_Q_/ν(CC)_Q_/ρ_b_(CH)_Q_], ∼1294 cm⁻¹ [ν(CC)_Q_/ρ_b_(CH)_Q_], ∼1204 cm⁻¹ [ν(ϕ)], ∼781 cm⁻¹ [ν(ϕ)], and ∼690 cm⁻¹ [ν_i.p._(Q)]. These bands indicate that the amide bond as well as the quinazoline and 3–chloro–4–fluoroaniline moieties participate in the interaction with the SB–AuNPs surface. Notably, no spectral features associated with PBA are detected, suggesting that when the compounds are introduced simultaneously or when Afatinib is adsorbed first, PBA molecules cannot effectively bind to the nanoparticle surface. This observation indicates a significantly stronger affinity of Afatinib toward SB–AuNPs compared with PBA. A different spectral pattern is observed when PBA is pre–adsorbed on the nanoparticles prior to the addition of Afatinib. In this case, the SERS spectrum displays bands characteristic of PBA at 1012 cm⁻¹ [ν(BO)] and 998 cm⁻¹ [δ_i.p._(ϕ)], together with a contribution to the band at 801 cm⁻¹ [δ(CCC), ν(BC), ν(OB)]. At the same time, features attributed to Afatinib remain visible at 1653 cm⁻¹, 1403 cm⁻¹, 801 cm⁻¹, and 783 cm⁻¹. This spectral response demonstrates that pre–adsorption of PBA enables the attachment of both molecules to the AuSBNP surface. In this configuration, PBA interacts with the nanoparticles through a perpendicular orientation of the phenyl ring, with the BO bond directed toward the surface, whereas Afatinib is anchored *via* the amide group together with the quinazoline and 3–chloro–4–fluoroaniline fragments. However, in all investigated systems, namely AuNPs + (Afatinib + PBA), AuNPs + Afatinib + PBA, and AuNPs + PBA + Afatinib, the interactions formed between the molecules and SB–AuNPs appear to be unstable. Figure S2 presents the time–dependent evolution of the spectral signals for each system at RT and 37 °C. At RT, desorption of the molecules from the AuNP surface is observed after 24 h. In contrast, at 37 °C, no characteristic spectral signal is detected even after 30 min following adsorption. These findings indicate that although Afatinib exhibits a relatively higher affinity toward AuNPs, the co–adsorption of PBA and this drug disrupts the adsorption efficiency. Consequently, both molecules desorb from the metal surface more rapidly than in approach where they are adsorbed individually.

## 4. CONCLUSIONS

In this study, AFM–SEIRA was applied for the first time to characterize the adsorption of Afatinib on 50 nm (HH–Au) and 25 nm (SB–Au) AuNPs. Complementary SERS measurements were used to assess the stability of the drug/AuNP interaction over time and under varying temperature conditions. The results can be summarized as follows:

1. Afatinib adsorbs on AuNP surfaces predominantly via the quinazoline moiety, with a significant contribution of the C=N and C–N bonds, indicating strong interaction of the heteroaromatic ring with the metal surface.
2. The amide group (–CONH–) plays a key role in binding, as evidenced by enhanced ν(C=O), ν(CN), and ρ_b_(NH) modes, confirming its direct involvement in molecule/metal interaction.
3. Amino groups (–NH–) contribute strongly to adsorption, with their role increasing over time, suggesting progressive molecular reorientation toward configurations favouring N–metal interactions.
4. The 3–chloro–4–fluoroaniline moiety interacts more weakly with the surface. The phenyl ring is located near the interface, while C–F and C–Cl substituents are oriented away from the metal.
5. Aliphatic fragments (CH₂ and CH₃ groups) exhibit low– and medium–intensity bands, indicating their proximity to the surface and auxiliary role in stabilizing the adsorption configuration.
6. The adsorption geometry depends on nanoparticle type, with stronger involvement of specific functional groups (e.g., C–N in quinazoline) observed for SB–AuNPs.
7. Time–dependent SERS measurements reveal that adsorption is dynamic, involving gradual reorientation of Afatinib molecules, with increasing contribution of amino groups and weakening of overall adsorption strength.
8. Desorption processes occur over time and are significantly accelerated at elevated temperature (37 °C), indicating limited stability of the drug/AuNP system under physiological conditions.

The co–adsorption studies of Afatinib and PBA revealed the following trends:

1. In competitive adsorption systems, Afatinib exhibits higher affinity to AuNPs than PBA, effectively dominating surface binding when co–adsorbed or adsorbed first.
2. Pre–adsorbed PBA enables temporary co–adsorption, however, interactions of both molecules are unstable, leading to faster desorption compared to single–component systems.

## Supporting Information

**Fig. S1.**
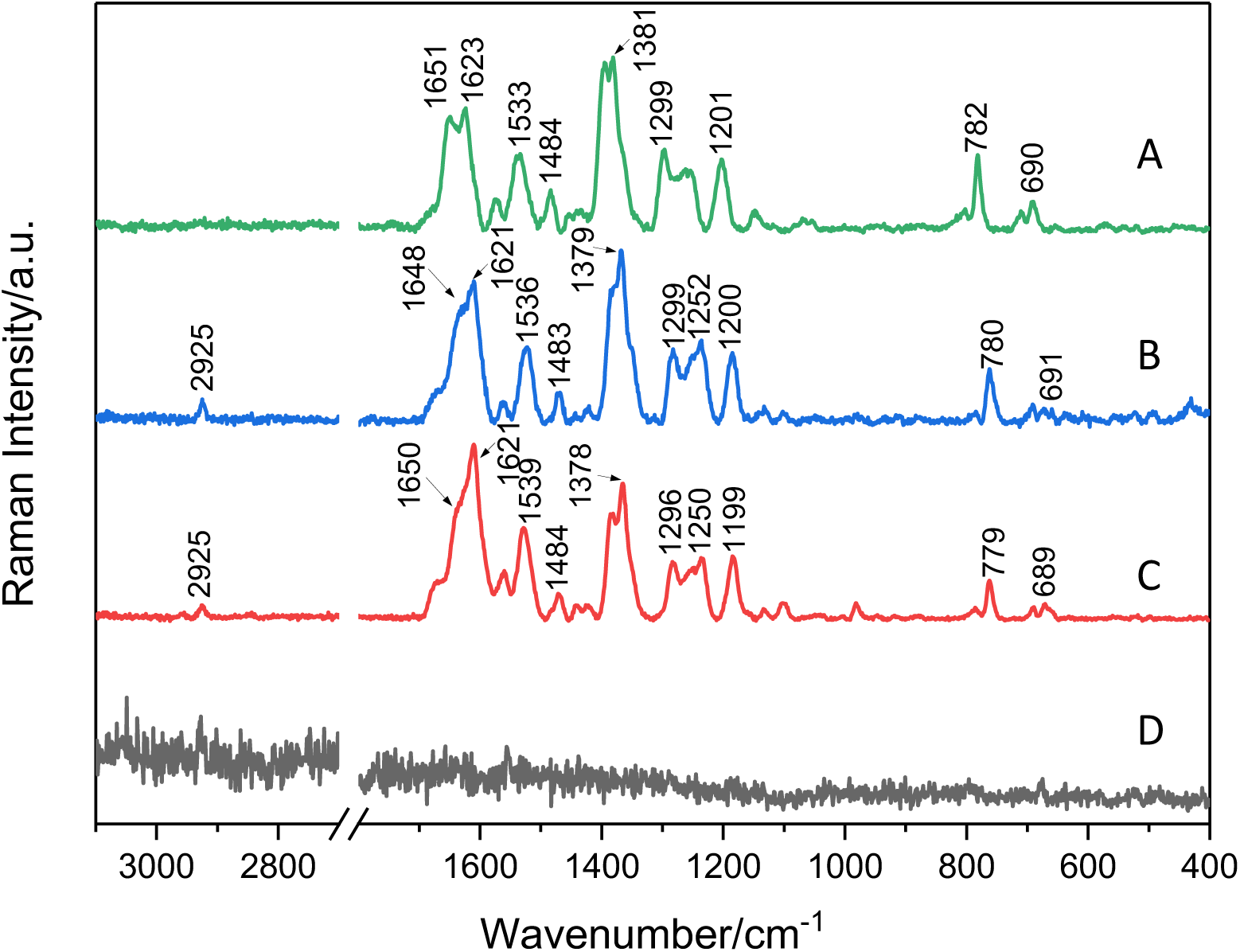
SERS spectra of Afatinib immobilized on the SB–AuNPs after 30 min (A), 4h (B), 8 h (C), and 24 h (D) adsorption time in the 37°C, respectively, in the spectral range of 3100–400cm^-1^.

**Fig. S2.**
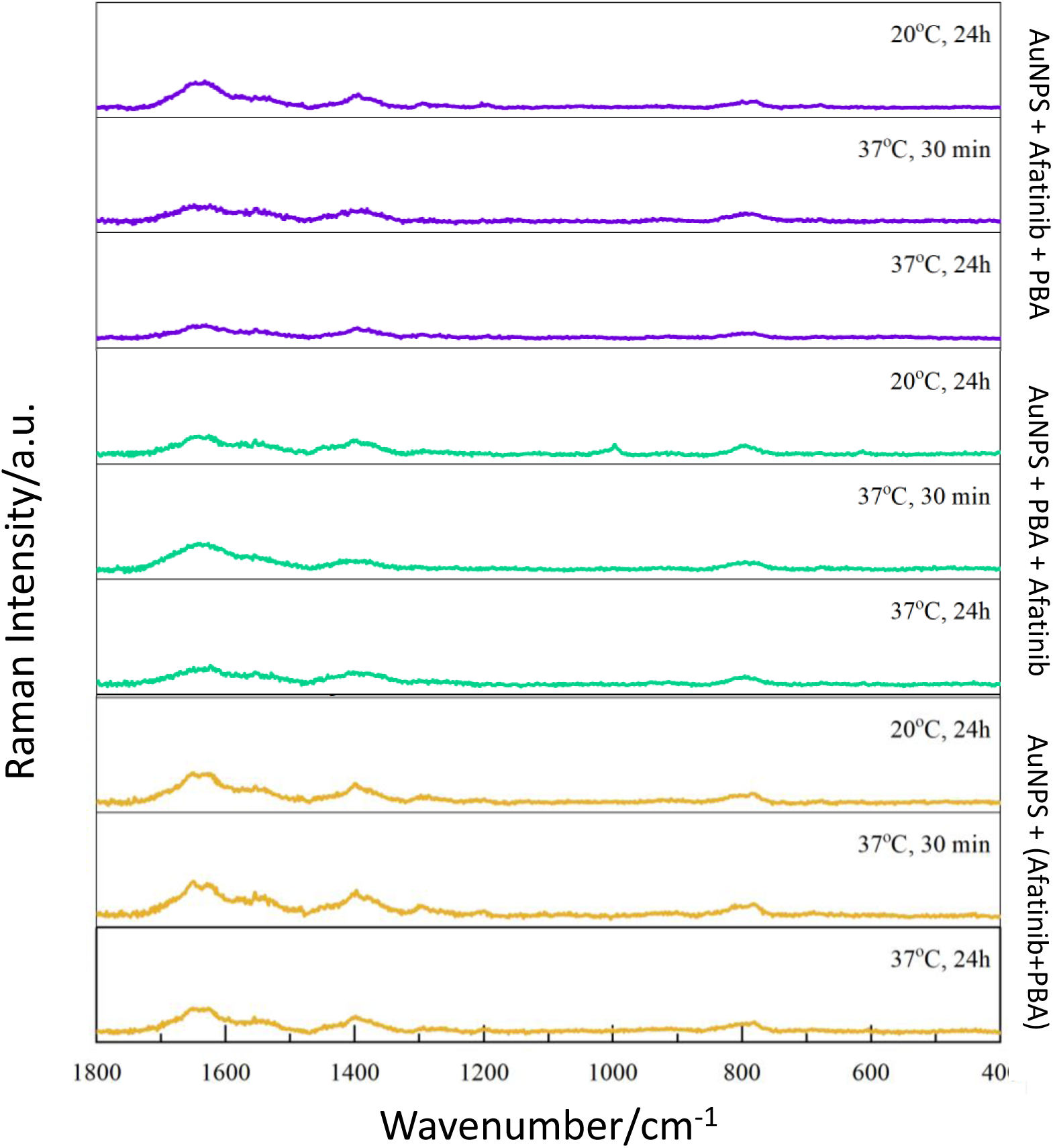
SERS spectra of three systems: AuNPs+Afatinib+PBA, AuNPs+PBA+Afatinib, and AuNPs+(Afatinib+PBA), recorded after 24 h of adsorption at RT and after 30 minutes and 24 h at 37°C, respectively, in the spectral range of 3100–400cm^-1^. Order of the listed components reflects the sequence of molecular attachment.

## Literature

[1] O. Afzal, A.S.A. Altamimi, M.S. Nadeem, S.I. Alzarea, W.H. Almalki, A. Tariq, B. Mubeen, B.N. Murtaza, S. Iftikhar, N. Riaz, I. Kazmi, Nanoparticles in Drug Delivery: From History to Therapeutic Applications, Nanomaterials 12 (2022) 1–27. 10.3390/nano12244494.

[2] I. Brigger, C. Dubernet, P. Couvreur, Nanoparticles in cancer therapy and diagnosis, Adv. Drug Deliv. Rev. 64 (2012) 24–36. 10.1016/j.addr.2012.09.006.

[3] N. Piergies, A. Dazzi, A. Deniset–Besseau, J. Mathurin, M. Oćwieja, C. Paluszkiewicz, W.M. Kwiatek, Nanoscale image of the drug/metal mono–layer interaction: Tapping AFM–IR investigations, Nano Res. 13 (2020) 1020–1028. 10.1007/s12274–020–2738–4.

[4] N. Piergies, M. Oćwieja, C. Paluszkiewicz, W.M. Kwiatek, Spectroscopic insights into the effect of pH, temperature, and stabilizer on erlotinib adsorption behavior onto Ag nanosurface, Spectrochim. Acta – Part A Mol. Biomol. Spectrosc. 228 (2020). 10.1016/j.saa.2019.117737.

[5] N. Piergies, J. Mathurin, A. Dazzi, A. Deniset–Besseau, M. Oćwieja, C. Paluszkiewicz, W.M. Kwiatek, IR nanospectroscopy to decipher drug/metal nanoparticle interactions: Towards a better understanding of the spectral signal enhancement and its distribution, Appl. Surf. Sci. 609 (2023). 10.1016/j.apsusc.2022.155217.

[6] N. Piergies, M. Oćwieja, M. Sadowska, D. Duraczyńska, M. Nattich–Rak, B.D. Napruszewska, AFM–SEIRA nanospectroscopy imaging of the drug adsorption on the PtNP monolayers, Meas. J. Int. Meas. Confed. 239 (2025). 10.1016/j.measurement.2024.115329.

[7] I. Brigger, C. Dubernet, P. Couvreur, Nanoparticles in cancer therapy and diagnosis, Adv. Drug Deliv. Rev. 64 (2012) 24–36. 10.1016/j.addr.2012.09.006.

[8] H. Cheng, J. Liao, Y. Ma, M. Tariq, Materials Today Bio Advances in targeted therapy for tumor with nanocarriers: A review, Mater. Today Bio 31 (2025) 101583. 10.1016/j.mtbio.2025.101583.

[9] J. Klekota, F.P. Roth, Chemical substructures that enrich for biological activity, Bioinformatics 24 (2008) 2518–2525. 10.1093/bioinformatics/btn479.

[10] M. Jelokhani–Niaraki, L.H. Kondejewski, L.C. Wheaton, R.S. Hodges, Effect of ring size on conformation and biological activity of cyclic cationic antimicrobial peptides, J. Med. Chem. 52 (2009) 2090–2097. 10.1021/jm801648n.

[11] García MA., Surface plasmons in metallic nanoparticles: fundamentals and applications., J. Phys. D Appl. Physics. 44 (2011) 283001.

[12] C. Noguez, Surface plasmons on metal nanoparticles: The influence of shape and physical environment, J. Phys. Chem. C 111 (2007) 3606–3619. 10.1021/jp066539m.

[13] M. Moskovits, Surface–enhanced spectroscopy, 1985.

[14] P.L. Stiles, J.A. Dieringer, N.C. Shah, R.P. Van Duyne, Surface–enhanced Raman spectroscopy, Annu. Rev. Anal. Chem. 1 (2008) 601–626. 10.1146/annurev.anchem.1.031207.112814.

[15] J.R. Lombardi, R.L. Birke, A unified view of surface–enhanced raman scattering, Acc. Chem. Res. 42 (2009) 734–742. 10.1021/ar800249y.

[16] M. Osawa, K.I. Ataka, K. Yoshii, Y. Nishikawa, Surface–enhanced infrared spectroscopy: The origin of the absorption enhancement and band selection rule in the infrared spectra of molecules adsorbed on fine metal particles, Appl. Spectrosc. 47 (1993) 1497–1502. 10.1366/0003702934067478.

[17] R.F. Aroca, R.E. Clavijo, M.D. Halls, H.B. Schlegel, Surface–enhanced raman spectra of phthalimide. Interpretation of the SERS spectra of the surface complex formed on silver islands and colloids, J. Phys. Chem. A 104 (2000) 9500–9505. 10.1021/jp002071q.

[18] M. Moskovits, Surface selection rules, J. Chem. Phys. 77 (1982) 4408–4416. 10.1063/1.444442.

[19] N. Piergies, M. Oćwieja, C. Paluszkiewicz, W.M. Kwiatek, Identification of erlotinib adsorption pattern onto silver nanoparticles: SERS studies, J. Raman Spectrosc. 49 (2018) 1265–1273. 10.1002/jrs.5384.

[20] N. Piergies, M. Oćwieja, C. Paluszkiewicz, W.M. Kwiatek, Applied Surface Science Nanoparticle stabilizer as a determining factor of the drug / gold surface interaction: SERS and AFM–SEIRA studies, Appl. Surf. Sci. 537 (2021) 147897. 10.1016/j.apsusc.2020.147897.

[21] P. Gnacek, N. Piergies, D. Duraczyńska, M. Kozak, C. Paluszkiewicz, M. Oćwieja, A physicochemical and spectroscopic characterization of novel erlotinib conjugates with platinum nanoparticles, Colloids Surfaces A Physicochem. Eng. Asp. 654 (2022). 10.1016/j.colsurfa.2022.130069.

[22] G. Palumbo, P. Jab, A. Golda, J. Koziel, B. Mingo, N. Piergies, D.L. Engelberg, Applied Surface Science Titanium coated with gold nanoparticles: a multifaceted investigation through electrochemical, spectroscopic, and biological approaches, 711 (2025). 10.1016/j.apsusc.2025.164032.

[23] P. Verma, Tip–Enhanced Raman Spectroscopy: Technique and Recent Advances, (2017). 10.1021/acs.chemrev.6b00821.

[24] A. Dazzi, F. Glotin, R. Carminati, Theory of infrared nanospectroscopy by photothermal induced resonance, J. Appl. Phys. 107 (2010). 10.1063/1.3429214.

[25] A. Dazzi, C.B. Prater, Q. Hu, D. Bruce, J.F. Rabolt, C. Marcott, focal point review AFM–IR: Combining Atomic Force Microscopy and Infrared Spectroscopy for Nanoscale Chemical Characterization, n.d.

[26] D. Święch, C. Paluszkiewicz, N. Piergies, E. Pięta, K. Kollbek, W.M. Kwiatek, Micro–and nanoscale spectroscopic investigations of threonine influence on the corrosion process of the modified fe surface by Cu nanoparticles, Materials (Basel). 13 (2020) 1–16. 10.3390/ma13204482.

[27] K. Kollbek, P. Jab, O. Magdalena, N. Piergies, Spectrochimica Acta Part A: Molecular and Biomolecular Spectroscopy Exploring the nanoscale: AFM–IR visualization of cysteine adsorption on gold nanoparticles, 318 (2024). 10.1016/j.saa.2024.124433.

[28] M. Schuler, J.C. Yang, K. Park, J. Kim, J. Bennouna, Y. Chen, C. Chouaid, non–small–cell lung cancer following chemotherapy, (2016) 417–423. 10.1093/annonc/mdv597.

[29] L. Zhang, Y. Luo, J. Chen, T. Cheng, H. Yang, C. Pan, H. Li, Z. Jiang, Efficacy and Safety of Afatinib in the Treatment of Advanced Non–Small–Cell Lung Cancer with EGFR Mutations: A Meta–Analysis of Real–World Evidence, 2021 (2021).

[30] X. Qian, L. Ge, K. Yuan, C. Li, X. Zhen, W. Cai, R. Cheng, X. Jiang, Targeting and microenvironment–improving of phenylboronic acid–decorated soy protein nanoparticles with different sizes to tumor, Theranostics 9 (2019) 7417–7430. 10.7150/thno.33470.

[31] T. Fu, Y. Wang, L. Wang, Y. Mao, ST6GALNAC1 mediates sialylation of mucins in non–small cell lung cancer to evade immune surveillance, 17 (2025) 6076–6089. 10.21037/jtd–2025–660.

[32] M. Oćwieja, Z. Adamczyk, M. Morga, A. Michna, High density silver nanoparticle monolayers produced by colloid self–assembly on polyelectrolyte supporting layers, J. Colloid Interface Sci. 364 (2011) 39–48. 10.1016/j.jcis.2011.07.059.

[33] M. Morga, Z. Adamczyk, Monolayers of cationic polyelectrolytes on mica – Electrokinetic studies, J. Colloid Interface Sci. 407 (2013) 196–204. 10.1016/j.jcis.2013.05.069.

[34] N. Piergies, M. Oćwieja, C. Paluszkiewicz, W.M. Kwiatek, Nanoparticle stabilizer as a determining factor of the drug/gold surface interaction: SERS and AFM–SEIRA studies, Appl. Surf. Sci. 537 (2021). 10.1016/j.apsusc.2020.147897.

[35] M.K. Trivedi, R.M. Tallapragada, A. Branton, D. Trivedi, G. Nayak, R.K. Mishra, Thermodynamics & Catalysis Biofield Energy Treatment: A Potential Strategy for Modulating Physical, Thermal and Spectral Properties of 3–Chloro–4–fluoroaniline, 6 (2015). 10.4172/2157–7544.1000151.

[36] A. Dwivedi, V. Baboo, A. Bajpai, Fukui Function Analysis and Optical, Electronic, and Vibrational Properties of Tetrahydrofuran and Its Derivatives: A Complete Quantum Chemical Study, 2015 (2015). 10.1155/2015/345234.

[37] N. Piergies, C. Paluszkiewicz, W.M. Kwiatek, Vibrational fingerprint of erlotinib: Ftir, rs, and DFT studies, J. Spectrosc. 2019 (2019). 10.1155/2019/9191328.

[38] M. Asakura, M. Okuno, Hyper–Raman Spectroscopic Investigation of Amide Bands of, (2021). 10.1021/acs.jpclett.1c01215.

[39] A. Manuscript, NIH Public Access, 138 (2014) 1665–1673. 10.1039/c2an36478f.Amide.

[40] P. Gnacek, N. Piergies, P. Niemiec, O. Kowalska, O. Magdalena, Spectrochimica Acta Part A: Molecular and Biomolecular Spectroscopy Spectroscopic studies under properties of chlorpromazine conjugated to gold nanoparticles, 320 (2024). 10.1016/j.saa.2024.124588.

[41] N. Piergies, E. Proniewicz, Y. Ozaki, Y. Kim, L.M. Proniewicz, In fl uence of Substituent Type and Position on the Adsorption Mechanism of Phenylboronic Acids: Infrared, Raman, and Surface– Enhanced Raman Spectroscopy Studies, (2013).

[42] N. Piergies, E. Proniewicz, Y. Kim, L.M. Proniewicz, Interaction of N–benzylamino(boronphenyl)methylphosphonic acid analogs with the gold colloidal surface under different concentration and pH conditions, in: J. Raman Spectrosc., John Wiley and Sons Ltd, 2014: pp. 581–590. 10.1002/jrs.4505.

[43] M. Kurt, An experimental and theoretical study of molecular structure and vibrational spectra of pentafluorophenylboronic acid molecule by density functional theory and ab initio Hartree Fock calculations, J. Mol. Struct. 874 (2008) 159–169. 10.1016/j.molstruc.2007.03.050.

44. [44] J Raman Spectroscopy – 2009 – Erdogdu – DFT FT-Raman FT-IR and NMR studies of 2-fluorophenylboronic acid.pdf, (n.d.).

